# Assessment of the Anticancer Activity of *Aegle marmelos* Leaf Extracts Against Ehrlich’s Ascites Carcinoma (EAC) Cell Line in *Swiss* Albino Mice

**DOI:** 10.1101/2025.03.23.641297

**Authors:** Md. Arafat Hossain, Md. Takdir Hossain

## Abstract

Cancer remains a significant health threat globally, and as a result, the demand for safer and more effective alternatives keeps growing. In this current study, we have assessed the anticancer activity of *Aegle marmelos* extract on Ehrlich Ascites Carcinoma (EAC) cells on Swiss Albino mice. Three extracts were prepared: a methanolic (APME), chloroform (APCE), and n-hexane (APHE) extract. These extracts were administered intraperitoneally in doses of 50 mg/kg and 100 mg/kg per dose on 6 consecutive days to mice. The anticancer activity was assessed from viable tumor cell count, with percentage (%) inhibition of cell growth, and changes in body weight of the mice that had been treated with extracts. Of all the extracts, APHE exhibited the most tumor growth inhibition (62.2% at 100 mg/kg), followed by APME (59.25%) and APCE (54.86%), compared to the standard chemotherapy drug doxorubicin (81.6% inhibition at 0.8 mg/kg/day). Mice treated with the extracts experienced less weight loss than the control group, with an approximate effect of 0.50 ± 0.06 g/day. A Brine Shrimp Lethality Bioassay demonstrated a cytotoxic dose-dependent impact, with LC50 values of 18.38, 36.79, and 46.45 micrograms per milliliter for n-hexane, methanolic, and chloroform extracts, respectively. The tumor suppression and cytotoxicity effects of *Aegle marmelos* extracts are probably attributable to the bioactive compounds such as flavonoids, alkaloids, and essential oils that have known apoptotic and antiproliferative activities. The extracts were not as effective as doxorubicin; however, the extracts natural composition and ability to reduce weight loss due to cancer indicate potential to be used in combination with or as an alternative to cancer-treating medications. Further research includes that mechanistic studies and clinical trials are needed to confirm their therapeutic potential.

## 1. Introduction

Cancer is a complicated disease that is driven by multilayered genetic and epigenetic alterations that lead to excessive growth and an increase in cellular populations (Singh & Roghini, 2023; Crispo et al., 2019). The development of the cancerous state is frequently associated with mutations in tumor suppressor genes (Weinberg, 1991), oncogenes (Croce, 2008), and microRNA genes (Frixa et al., 2015). As a major public health concern, cancer remains a significant challenge in medical research and treatment (Wong, 2011; Fadeel & Orrenius, 2005). Conventional treatments such as hormone therapy (Abraham & Staffurth, 2016), radiation (Allen et al., 2017; Schaue & McBride, 2015), chemotherapy (Nygren, 2001; Mitchison, 1979), and surgery (Tohme et al., 2017; Wyld et al., 2015), are often used but frequently have major adverse effects. The high death rate due to cancer, as well as the side effects of anticancer drugs, have led researchers to search for new therapeutic options with better efficacy and lower toxicity (Ioele et al., 2022). These difficulties resulted in researchers’ pursuit of natural products that may have anticancer activity (Cragg et al., 2009; Lichota & Gwozdzinski, 2018). There are many plant-derived bioactive compounds, such as terpenoids, phenolic acids, flavonoids, tannins, lignans, quinones, coumarins, and alkaloids with potential antioxidant activity, which may be helpful in preventing or treating cancer (Hilal et al., 2024). It has been reported that antioxidants exert antistress, antitumor, antimutagenic, and anticarcinogenic effects, which inhibit the growth of cancer cells and enhance immune function (Mohamed et al., 2017; Katiyar, 2005; Roleira et al., 2015).

Despite advancements in modern medical science, Ayurvedic treatments continue to be valued for their therapeutic benefits (Aggarwal et al., 2006). Consequently, Recent research has, therefore, explored plant-based bioactive compounds with potential anticancer activity (Shrihastini et al., 2021). *Aegle marmelos* (figure 1), also called Bael, Bengal quince, wood apple, or stone apple, is a medicinal tree belonging to the Rutaceae family (Kohli et al., 2022; Baliga et al., 2013). Indigenous to the Indian subcontinent and Southeast Asia, *Aegle marmelos* is widely regarded for its cultural and medicinal significance, particularly in Ayurveda (Vasava et al., 2018). This tropical tree bears fruit with well-documented medicinal properties and is cultivated extensively across Asia. Its fruit, leaves, and bark contain bioactive compounds such as flavonoids, alkaloids, and essential oils, which contribute to its therapeutic potential. Traditionally, different parts of the plant have been utilized as digestive aids, astringents, antidiabetic agents, and antimicrobial agents (Venthodika et al., 2021). It has even been used to treat gastrointestinal disorders (Rao et al., 2003) and respiratory diseases, including asthma and bronchitis (Ghatule et al., 2014; Timalsina et al., 2021), as well as chronic diseases, including diabetes, hypertension, arthritis, and skin infections (Kohli et al., 2022). Recent scientific studies highlight *Aegle marmelos* for its antioxidant (Sabu & Kuttan, 2004) and anticancer potential (Gupta et al., 2016), making it a promising candidate for herbal-based cancer treatments. Given its bioactive composition and therapeutic properties, the present study aims to explore the anticancer potential of *Aegle marmelos* leaf extracts using an *in vivo* mouse model.

**Figure 1.**
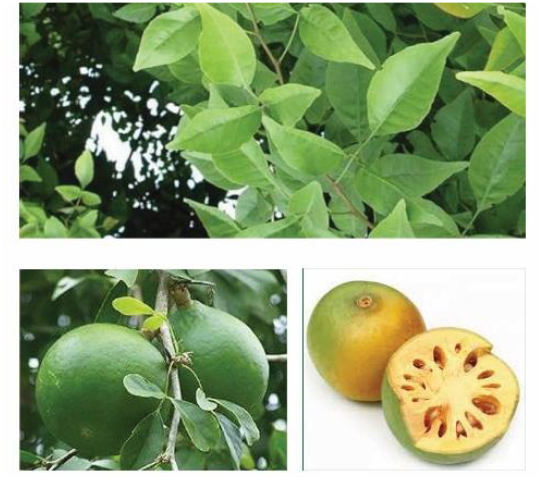
Botanical Image of *Aegle marmelos*

### Taxonomy

Domain:Eukaryota Kingdom: Plantae

Phylum: Spermatophyta

Subphylum:Angiospermae Class: Dicotyledonae

Subclass: Rosidae

Order:Sapindales Family:Rutaceae Genus: *Aegle*

Species: *Aegle marmelos*

## 2. Phytochemistry of *Aegle marmelos*

*Aegle marmelos* is a known safe reservoir of many phytoconstituents, and the most important include marmin, marmenol, marmelosin, marmelide, rutaretin, psoralen, alloimperatorin, scopoletin, marmelin, aegelin, fagarine, anhydromarmelin, β-phellandrene, limonene, betulinic acid, imperatorin, marmesin,luvangentin, marmelosin, and auroptene (Bansal & Bansal, 2011). For example, according to Yadav et al, the quantity of tannin and riboflavin in *Aegle marmelos* was identified as 0.985% and 0.005% respectively (Yadav et al. 2011). Additionally, Bansal & Bansal successfully separated and measured several organic acids, such as oxalic, malic, tartaric, and ascorbic acids, using reverse-phase high-performance liquid chromatography (RP-HPLC) (Bansal & Bansal, 2011). Prakash et al. used analytical methods such as LC-MS, LC-MS/MS, and HPLC to identify several phenolic compounds in the fruit, including chlorogenic acid (136.8 µg/g), ellagic acid (248.5 µg/g), ferulic acid (98.3 µg/g), gallic acid (873.6 µg/g), protocatechuic acid (47.9 µg/g), and quercetin (56.9 µg/g) (Prakash et al., 2011). Charoensiddhi et al. conducted a study using SPME (solid phase micro-extraction) coupled with GC/MS to detect volatile components of *Aegle marmelos*. They detected a wide variety of constituents, including hexanal, limonene, isoamyl acetate, β-phellandrene, acetoin, p-cymene, (E, E)-2,4-heptadienal, I-2-octenal, dehydro-p-cymene, 3,5-octadiene-2-one, linalool oxide, α-Cubebene, citronellal, trans-p-mentha-2,8-dienol, β-cubebene, β-caryophyllene, hexadecane, pulegone, verbenone, α-Humulene, carvone, carvyl acetate, carvone, dihydro-β-Ionone, β-Ionone, I-6,10-dimethyl-5,9-undecadien-2-one, humulene oxide, caryophyllene oxide and _exadecenoic acid examining *Aegle marmelos* oil. (Charoensiddhi et al., 2008). Moreover, the seed oil has been reported to contain common fatty acids, including oleic, stearic, linoleic, palmitic, and linolenic acids (Dhankhar et al., 2011). Interestingly, Katagi et al. identified ricinoleic acid, an unusual fatty acid constituting approximately 12.5% of the total fatty acid content in the seed oil, along with the more commonly occurring fatty acids (Katagi et al. 2011).

## 3. Materials and Methods

### 3.1 Chemicals and reagents

All chemicals and reagents utilized in this study were analytical grade and purchased from reputable suppliers. Doxorubicin was acquired from Beacon Pharmaceutical Limited, Bangladesh, and used as the standard chemotherapeutic agent. The drug was prepared in distilled water at a concentration of 0.075 mg/ml for intraperitoneal injection. Methanol, chloroform, and n-hexane (HPLC grade) were purchased from Sigma-Aldrich (USA) for use in leaf extracts. Dimethyl sulfoxide (DMSO) (Merck, Germany) was used at a 2% concentration as a solvent for dissolving the extracts in experimental treatments. Trypan blue dye (0.4%) was utilized for cell viability assays to differentiate between viable and non-viable EAC cells. Normal saline solution (0.98% NaCl) from Beximco Pharmaceuticals Ltd, Bangladesh, was prepared for diluting tumor cell suspensions before transplantation. Cotton and Whatman No. 1 filter paper were used for extract filtration, while phosphate-buffered saline (PBS) was employed for cell washing and other experimental procedures. All solutions were freshly prepared before use and stored under appropriate conditions to maintain stability and effectiveness.

### 3.2 Extract preparation

The plant *Aegle marmelos* was collected from a plain local site in Patuakhali, Bangladesh, and was identified by the Bangladesh National Herbarium, Dhaka (Accession Number: 58997). The extraction method was carried out following maceration techniques as outlined by Azwanida (2015) and Sasidharan et al. (2011). The solvents used for extraction were methanol, chloroform, and n-hexane. *The leaves were shade-dried to preserve bioactive compounds, following recommendations by Harborne (1998).* To remove dirt, the collected fresh leaves of the *Aegle marmelos* were washed with cold water and then hot water to remove any kind of organism. The leaves were dried in a shaded area for 15 days (at room temperature, 25 ± 2 °C) to prevent photo-oxidation of bioactive components and then ground into a fine powder. The leaf’s fine powder (500 g) was extracted by using APME, APCE, and APHE at room temperature (25 ± 2 °C) for 15 days with agitation three times a day. Finally, filtration was performed by using cotton and then No. 1 Whatman filter paper. The filtrate is then concentrated by using a rotary evaporator and finally air-dried. The extract yield was 6.78% (w/w) with methanol, 7 % (w/w) with chloroform, and 7.8% (w/w) with n-hexane.

### 3.2 Experimental animal

Swiss albino mice, 5-7 weeks of age and weighing 20-26 grams, were acquired from the Department of Pharmacy, Jahangirnagar University, Dhaka, Bangladesh. The mice were housed in standard environmental conditions in the Molecular Biology lab, having a relative humidity (RH) of 55 ± 5% and a temperature of 22 ± 2 °C. Before the experiment, the mice were acclimated for one week in the animal house in a 12-hour light/12-hour dark cycle. The mice have an approximate life span of 2-3 years, with a possible lifespan up to 4 years. They were kept in iron cages on sawdust and straw bedding, which was changed weekly. Mice were provided a standard diet prepared by ICDDR’B and fresh water. The animal house was maintained at room temperature between 25-32 degrees Celsius under a 14-hour light and 10-hour dark cycle. The pellet diet procured from ICDDR’B in Dhaka was as follows per 100 grams of diet: starch 66 grams, casein 20 grams, fat 8 grams, vitamins – standard vitamins-2 grams, and salt 4 grams.

### 3.4 Ehrlich’s ascites carcinoma (EAC) cell line

Experimental tumors are indispensable models for cancer research, providing insights into tumor growth, progression, and response to treatment. EAC is one of the most frequently used tumor models. EAC cell lines are used in cancer studies globally due to their rapid proliferation, plasticity, and capacity to reflect aspects of tumor behaviour. The tumor was initially discovered by the famous scientist Paul Ehrlich in 1907 when he discovered the EAC tumor in the mammary gland of a white mouse. This was the start of research on experimental tumors. Later, there was the development with regard to the present form of EAC cells by researchers Loewenthal & John, where they derived EAC from other mammary gland-originated cell lines, improving the model for scientific use.

EAC cells have a thin, but solid, outer membrane with a very solid membrane matrix. Studies on the morphology of normal and cancer cells have shown that intracellular and surface membranes conform to the ‘unit membrane’ structure. This fundamental structure is a bimolecular lipid leaflet, and on each side of its phospholipid bilayer, some proteins or polysaccharides are important for maintaining the integrity and function of the cell. The EAC cells, like other cancer cells, have this unit membrane organization, and they continue to do so, and that is why they are rapidly proliferating and becoming resistant to drug treatment.

EAC tumors can be grown in experimental studies in two forms: either solid or ascitic. The solid form is grown subcutaneously, and the ascitic form is grown using injecting a tumor cell suspension directly into the peritoneal cavity of a mouse. The ascitic tumor is observed as a milky white fluid, filled with large, round tumor cells that reproduce very quickly. One million EAC tumor cells can expand to 25-100 million cells/mL in days. Mice bearing these tumors typically survive for 14-30 days, depending on tumor burden and host resistance. This short survival time makes EAC a useful model system for the study of tumor growth patterns, drug response, and interactions with the immune system in cancer research.

### 3.5 Microscope

We viewed the samples loaded on the hemocytometer using a binocular microscope set to magnification levels of 10x, 40x, and 100x for viewing. At the 10x level of magnification, low power was used for a general view of the distribution of cells throughout the grid on the hemocytometer. This allowed us to take note of areas of interest as well as get a rough density count of the cells. At the 40x magnification level, more structural cellular properties could be distinguished, making it easier to discern the cellular forms and arrangements that are important for comparison between various types of cells. Lastly, the 100x magnification level exposed high detail, which demonstrated subtle morphological properties that were necessary to confirm cell identification. This methodical approach enabled us to efficiently confirm cell identifications, count them accurately, and perform a comprehensive analysis of the sample.

### 3.6 Hemocytometer

The hemocytometer is one of the most widely used, cost-effective, and accurate tools for determining the number of cells in samples. This is an inexpensive yet specialized device perfect for microscopic counting of cells and provides an accurate assessment of the cell concentration in many different biological and medical applications (Chen & Chiang, 2024). A hemocytometer consists of a thick glass slide with two designated counting chambers, each systematically divided into nine large squares, each measuring 1 mm^2^in area. These chambers are on a surface etched and silvered, separated by a central groove used to reinforce the structural integrity of the slide and become a means for easy loading of the cell suspension (Chen et al., 2024). To ensure accurate cell counting, a coverslip is placed on top of the raised supports of the ‘H’-shaped troughs, enclosing both counting chambers and ensuring a uniform sample distribution. Each counting chamber is further equipped with a ‘V-shaped notch or an indentation at either end, where the prepared cell suspension is carefully introduced into the chamber. After the cell suspension is loaded, it is equally distributed throughout the chamber and occupies a specific, defined volume of liquid. The counting chamber has an engraved grid on the top that provides an organized and systematic way to count the cells without making errors or discrepancies. Next, the hemocytometer was placed on the stage of the microscope, and the cell suspension was viewed at appropriate magnification for counting (Absher, 1973). By counting the number of cells in the designated grid areas and multiplying by the appropriate dilution factors, researchers can determine the cell concentration per unit volume. The cell concentration per ml was determined using the following process.

Number of cells per ml

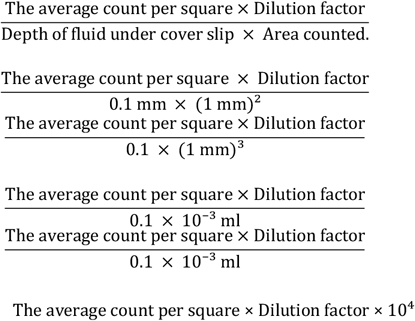

### 3.7 Preparation of stock solution for the test samples

The preparation of stock solutions for the test samples involved dissolving *Aegle marmelos*leaf extracts in a solvent to ensure proper administration and bioavailability in the experimental animals. Three extracts—APME, APCE, and APHE—were prepared using 2% DMSO, selected for its ability to dissolve a wide range of organic compounds and its compatibility with biological systems at low concentrations (Modrzyński et al., 2019). Each extract was dissolved in DMSO to create a stock solution with a concentration of 100 mg/mL, which was then diluted with phosphate-buffered saline (PBS) to achieve working concentrations of 50 mg/kg and 100 mg/kg for intraperitoneal (i.p.) administration. For instance, a 20-gram mouse received 0.1 mL for a 50 mg/kg dose and 0.2 mL for a 100 mg/kg dose. In addition, doxorubicin, used as the standard chemotherapeutic agent, was dissolved in distilled water at 0.075 mg/mL and administered at a dose of 0.8 mg/kg/day i.p. The control group was given a solution containing 2% DMSO in PBS, ensuring that any observed effects were attributable to the extracts rather than the solvent. All stock solutions were freshly prepared before each administration to maintain stability and efficacy, stored at 4°C when not in use, and brought to room temperature before administration, thereby ensuring uniform dosing and accurate assessment of the anticancer effects of the extracts.

### 3.8 Experimental tumor model

The EAC cell line used in this study was initially obtained from the Department of Biochemistry and Molecular Biology, University of Rajshahi, Bangladesh. To ensure a continuous and viable supply of tumor cells for experimental use, the culture and maintenance of EAC cells were carried out following the established protocol described by Alam et al. (2016), with slight modifications tailored to the specific laboratory conditions. For propagation, EAC cells were maintained in *Swiss*Albino mice through serial intraperitoneal (i.p.) transplantation. Every week, freshly isolated EAC cells were collected from a donor mouse that had been inoculated with tumor cells six to seven days prior. The ascitic fluid containing tumor cells was carefully withdrawn using a sterile syringe and subsequently diluted with normal saline solution (1% NaCl) to ensure an optimal cell concentration for transplantation. To assess cell viability and facilitate the separation of macrophages from tumor cells, the diluted cell suspension was placed in a sterile petri dish and incubated at 37.5°C in an air incubator for one hour.

During the incubation period, macrophages adhered firmly to the bottom surface of the culture vessel, allowing them to be distinguished from the free-floating tumor cells. This selective adherence facilitated the effective separation of macrophages from the EAC cells, ensuring a purer tumor cell suspension for subsequent experiments. After incubation, the petri dish was gently agitated using a brief vortexing step to resuspend any loosely attached tumor cells. The supernatant, now enriched with EAC cells, was carefully collected using a sterile pipette and transferred to a fresh container for further processing. To standardize the tumor cell concentration for experimental use, the collected suspension was carefully adjusted to approximately 2 × 10^6^cells/ml. This was achieved using a hemocytometer, a specialized counting chamber that provides an accurate measurement of cell density. To ensure the viability of the tumor cells, a 0.4% trypan blue dye exclusion assay was performed. Throughout the transplantation process, strict aseptic conditions were maintained to ensure accuracy and reproducibility.

### 3.9 Transplantation of ascitic tumor

Ascitic fluid was drawn out from different tumor-bearing *Swiss*albino mice at the respective log phases of tumor cells. A 3ml syringe filled with a 20-gauge needle was used for this tumor cell aspiration. The freshly drawn fluid was diluted with normal saline (0.98% NaCl solution), and the tumor cell number was adjusted to approximately 2106 cells/ml by counting the number with the help of a hemocytometer. The viability of tumor cells was observed by the trypan blue dye (0.4%) exclusion assay. Cell samples showing above 90% viability were used for transplantation. A tumor suspension of 0.1 ml was injected intraperitoneally (i.p.) into each Swiss albino mouse. The strict aseptic condition was maintained throughout the transplantation process. The intraperitoneal route of administration, along with careful attention to dosing volume and needle selection, was chosen based on established laboratory animal handling protocols, as supported by Turner et al. (2011). *In vivo,*the antitumor activity of the test samples was determined by measuring the effect of the samples on the tumor cell growth of EAC cell-bearing mice.

### 3.10 Studies on EAC Cell Growth Inhibition

To determine the cell growth inhibition of the leaf extracts, five groups of *Swiss*albino mice (6 in each group) weighing 20-25 g were used. For therapeutic evaluation, 136×10_4_EAC cells in every mouse were inoculated into each group of mice on day 0. Treatments were started after 24 hours of tumor inoculation and continued for six days. Groups one to three or every APME, APCE, and APHE received the test compound at the doses of 50 mg/kg and 100mg/kg/day (i.p.). The standard group received Doxorubicin at the dose of 0.8 mg/kg/day (i.p.), and group five was used as a control. On day six, the mice in each group were sacrificed, and the total intraperitoneal tumor cells were collected using 0.98% normal saline. Viable cells were first identified by using trypan blue and then counted by a hemocytometer. The total number of viable cells in every animal of the treated groups was compared with that of the control (EAC treated only).

### 3.11 Evaluation of Weight Loss

To assess weight loss, the weight of each mouse was measured daily for six consecutive days following the inoculation of carcinoma cells, using an electronic balance (KAMRY, electronic kitchen scale, China) during the treatment period. The rate of weight loss was then calculated by taking the average weight values for both the control and treated mice, using the following equation:

The rate of weight loss

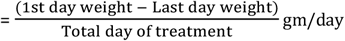

### 3.12 Brine Shrimp Toxicity Assessment

The Brine Shrimp Toxicity Bioassay was conducted following a conventional method with minor modifications. *Artemia salina*nauplii were used as the test organism for this study. For the experiment, 16 mg of the plant extract was dissolved in 250 μL of pure DMSO and diluted to a final volume of 25 mL with seawater, obtaining a stock solution with a concentration of 900 μg/mL. Serial dilutions were performed to prepare solutions of varying concentrations (200, 150, 100, 75, 50, 25, 12.5, and 6.25 μg/mL). Each pre-marked vial contained 2.5 mL of seawater, to which 2.5 mL of the respective plant extract solution was added, making a final volume of 5 mL. Ten live brine shrimp nauplii were introduced into each vial. The experiment was conducted twice, and after 24 hours, the number of surviving nauplii was counted using a magnifying glass and recorded for analysis.

## 4. Results

### 4.1 EAC cell growth inhibition

The *in vivo*antitumor activity of different extracts—APME, APHE, and APCE—from *Aegle marmelos*leaves was evaluated in EAC cell-bearing mice. The assessment was based on viable EAC cell count and percentage inhibition of cell growth. In the untreated EAC control group, the average number of viable tumor cells per mouse was recorded as (10.12 ± 2.72) × 10^7^cells/ml. Treatment with Doxorubicin (0.8 mg/kg/day) resulted in the most significant reduction in viable cells, with an EAC cell growth inhibition of 81.6%, showing a viable cell count of (1.87 ± 0.05) × 10^7^cells/ml.

Treatment with APME at doses of 50 mg/kg and 100 mg/kg significantly reduced the viable cell count (P<0.05). APME exhibited 50.31% and 59.25% EAC cell growth inhibition at 50 mg/kg and 100 mg/kg, respectively, with viable cell counts of (3.95 ± 1.87) and (3.20 ± 2.06) × 10^7^cells/ml. Similarly, APCE demonstrated 30.21% and 54.86% inhibition at 50 mg/kg and 100 mg/kg, respectively, with viable cell counts of (6.2 ± 2.25) and (3.90 ± 1.56)× 10^7^cells/ml. In comparison, APHE showed the highest inhibition among the extracts, with 50.60% and 62.2% EAC cell growth inhibition at 50 mg/kg and 100 mg/kg, respectively, with viable cell counts of (3.68 ± 1.80) and (2.3 ± 1.34) × 10^7^cells/ml.

The order of EAC cell growth inhibition by different treatments was as follows: Doxorubicin > APHE > APME > APCE. The results of growth inhibition are presented in Table 1, and the graphical representation is shown in Figure 2.

**Table 1.**
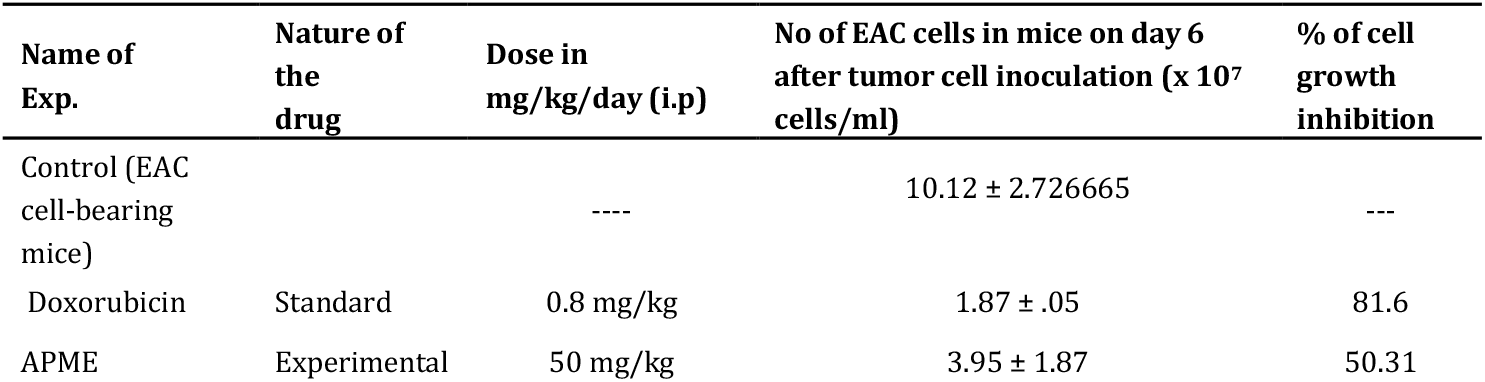
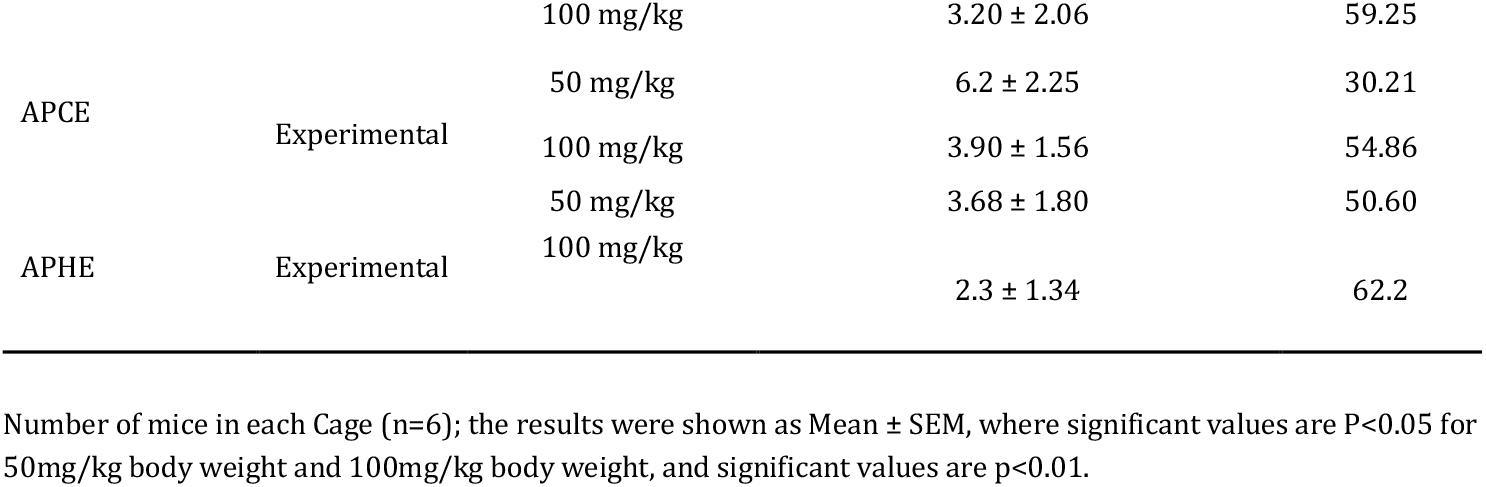
Effect of *Aegle marmelos*leaf extracts on Ehrlich ascites carcinoma (EAC) cell growth

**Figure 2.**
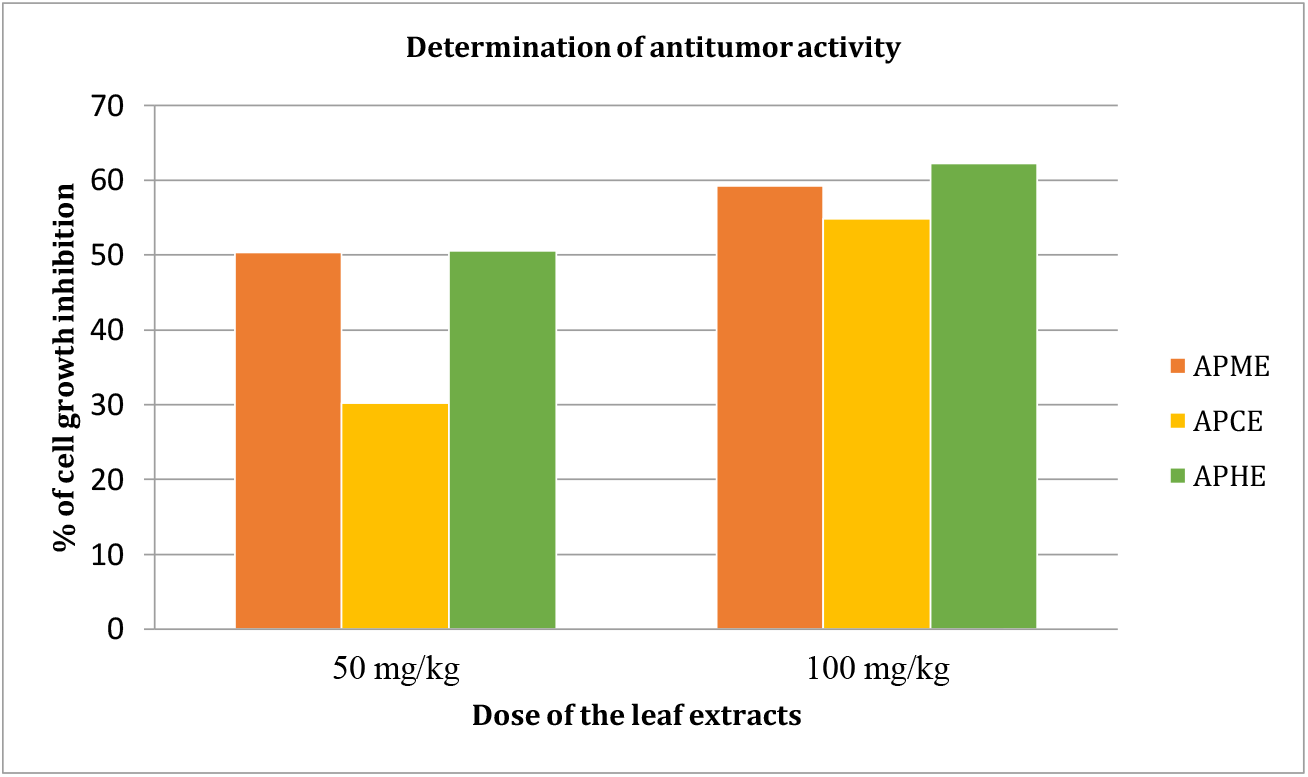
Determination of Antitumor Activity of *Aegle marmelos*Leaf Extracts. This figure presents the percentage inhibition of EAC cell growth in *Swiss*Albino mice treated with different *Aegle marmelos*leaf extracts (APME, APCE, and APHE) at 50 mg/kg and 100 mg/kg doses. The results are compared to the standard chemotherapeutic drug doxorubicin (0.8 mg/kg/day). Among the extracts, APHE exhibited the highest tumor growth inhibition (62.2% at 100 mg/kg), followed by APME and APCE, while doxorubicin showed the highest inhibition (81.6%). The figure visually represents the efficacy of these extracts in suppressing tumor growth.

Number of mice in each Cage (n=6); the results were shown as Mean ± SEM, where significant values are P<0.05 for 50mg/kg body weight and 100mg/kg body weight, and significant values are p<0.01.

### 4.2 Evaluation of Weight Loss

Weight loss is a common symptom of cancer. In the experiment, the mice gradually lost weight after the inoculation of cancer cells. The rate of weight loss was 1.3 ± 0.08 g/day for the control mice and approximately 0.50 ± 0.06 g/day for the leaf-treated mice. The leaf-treated mice showed a lower rate of weight loss compared to the control mice exhibited the highest rate of weight loss, as depicted in Figure 3.

**Figure 3.**
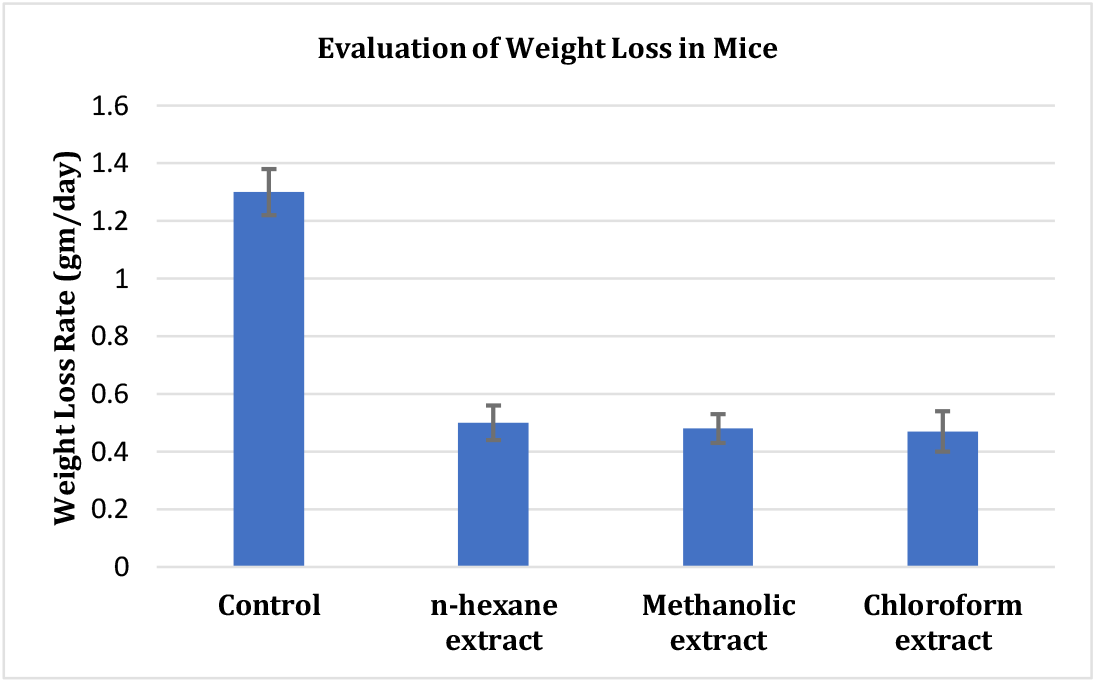
Weight loss evaluation in *Swiss*Albino mice following treatment with *Aegle marmelos*leaf extracts. Data are presented as mean ± SD (n = 6) for all tested doses. Significant differences from the control group are indicated, with statistical significance levels set at p < 0.05 and p < 0.001.

### 4.3 Brine Shrimp Toxicity Assessment

The cytotoxic potential of methanolic, n-hexane, and chloroform extracts of *Aegle marmelos*leaves was assessed using the brine shrimp lethality assay. The results revealed significant toxicity in a dose-dependent manner. At a concentration of 70 µg/mL, the methanolic extract induced 100% mortality. The LC50 values, calculated from the best-fit line slope, were 18.38, 36.79, and 46.45 µg/mL for n-hexane, methanolic, and chloroform extracts, respectively. The pronounced lethality of *Aegle marmelos*leaf extracts indicates the presence of potent cytotoxic compounds, highlighting the need for further investigation into their potential anticancer properties.

## 5. Discussion

This research was conducted to investigate the anticancer effect of *Aegle marmelos*(Bael) leaf extracts: APME, APCE, and APHE against EAC in Swiss Albino mice. This research aimed to assess the efficiency of *Aegle marmelos*extracts on preventing tumor cells from proliferating, reducing cancer-induced cachexia, and inducing cytotoxicity. The proposed mode of action is likely multi-targeted: modulation of oxidative stress, induction of apoptosis, arresting the cell cycle, and inhibition of tumor-related inflammation. These effects are potentially mediated by various phytochemicals, including marmelin (HDNC), lupeol, eugenol, citral, and imperatorin (Sudhamani et al., 2014; Neha et al., 2021; Sushmitha et al., 2021; Venkatakarthikeswari et al., 2021), which have displayed antiproliferative, antioxidant, and pro-apoptotic activities in multiple cancer models.

The findings revealed that all extracts inhibited EAC cell growth significantly In a dose-dependent manner, with APHE at 100 mg/kg showing the highest inhibition (62.2%), followed closely by APME (59.25%) and APCE (54.86%). These values, although not as high as doxorubicin (81.6%), indicate substantial antitumor activity. Notably, APHE had the lowest LC_50_ value (18.38 µg/mL) in the brine shrimp lethality assay, further supporting its cytotoxic strength. Mice treated with these extracts also showed lower weight loss rates (0.50 ± 0.06 g/day), suggesting that the extracts may alleviate cancer-associated cachexia. This effect could be linked to the anti-inflammatory and immunomodulatory actions of flavonoids, which have been shown to preserve muscle mass and improve metabolic function during cancer progression (Mahato & Kumar,2022; Pynam & Dharmesh, 2018; Li et al., 2022).

Comparatively, these observations are consistent with many earlier studies that documented the anticancer effects of *Aegle marmelos*. Gupta et al. documented tumor suppression in DMBA-induced skin carcinogenesis that was associated with the bark extract and indicated further reliance on ROS-mediated apoptosis (Gupta et al. 2016). Baliga et al. investigated these anticancer effects of *Aegle marmelos*using fruit pulp and demonstrated the loss of integrity in the mitochondrial membrane as well as caspase-3 activity (Baliga et al., 2012). Sabu and Kuttan showed that their diabetic model, which showed antioxidant activity, was the result of lower lipid peroxidation and increased superoxide dismutase (SOD) level (Sabu and Kuttan, 2004). These results support our findings and confirm the presence of bioactive principles capable of regulating oxidative stress—a central driver in cancer pathogenesis.

*Aegle marmelos*contains various phytoconstituents that demonstrate anticancer mechanisms. One of these constituents, apigenin, a flavonoid frequently reported on *Aegle marmelos*, binds to Bcl-2 proteins with increased activity of Bax, inducing cytochrome c release and caspase activation (Imran et al., 2020). Luteolin, another phytoconstituent, inhibits phosphorylation of AKT and ERK, induces G1 cell cycle arrest, and downregulates cyclin D1 expression (Ong et al., 2008). *Aegle marmelos*alkaloids can affect tubulin similarly to vinblastine and paclitaxel, disrupting microtubule formation (Wang et al., 2016). Essential oils such as limonene and eugenol present in bael are also linked to anti-angiogenic and apoptosis-inducing effects (Padhy et al., 2016; Venthodika et al., 2020). These overlapping mechanisms suggest that the observed antitumor activity is not due to a single agent, but to a synergistic phytochemical network—a hallmark of botanical therapeutics. Beyond *Aegle marmelos*, the Rutaceae family contains members that exhibit anti-cancer potential. *Ruta graveolens*, another Rutaceae member, reported cytotoxicity in colon and breast cancer forms via DNA intercalation and topoisomerase inhibition (Arora et al., 2021; Fadlalla et al., 2011). This suggests that not only are the anticancer traits in *Aegle marmelos*, but also potential evolutionarily conserved traits in Rutaceae species, in addition to exploring the bioprospecting of related species.

While these findings are promising, there are limitations to consider. Firstly, the efficacy of the extracts, though significant, did not reach the inhibitory levels of doxorubicin. Secondly, this study did not perform molecular profiling of apoptosis (e.g., expression of p53, Bcl-2, caspases) or conduct histopathological analysis of tumor tissues. Thirdly, chronic toxicity assessments were not conducted, which is essential before proposing these extracts for long-term or human use. Furthermore, the EAC model, being a rapidly growing transplantable tumor, may not fully mimic the complexity of human malignancies. Hence, further investigations using solid tumor models, xenografts, and clinical samples are needed.

In summary, this study demonstrates that leaf extracts of *Aegle marmelos*, especially the n-hexane fraction (APHE), have anti-cancer activity in vivo and may additionally aid in reducing cancer-related weight loss. The findings support the use of *Aegle marmelos*as supportive care in cancer treatment and provide a rationale for future studies. Mechanistic studies involving gene expression studies, receptor binding, and metabolomic approaches, in conjunction with preliminary toxicological and pharmacokinetic studies, will be pivotal in converting the findings into clinical applications or phytopharmaceutical preparations.

## 6. Conclusion

This study provides evidence for the anticancer potential of *Aegle marmelos*leaf extract against EAC in Swiss Albino mice. The APHE exhibited the highest inhibition of tumor growth (62.2% at a concentration of 100 mg/kg) when compared with the other extracts, APME and APCE. The extract-treated groups also showed less weight loss, indicating their potential influence on cancer cachexia. Although the extracts displayed lower efficacy compared to the standard chemotherapeutic agent doxorubicin (81.6% inhibition), their natural origin and lower toxicity highlight their potential as safer alternatives. Certain bioactive compounds, including flavonoids, alkaloids, and essential oils, can probably contribute to the anticancer effects via apoptosis induction and tumor growth inhibition. Even though these indications are promising, additional studies are required to determine the exact molecular mechanisms, dosage optimization, and long-term safety. Clinical trials will be needed to confirm the therapeutic properties of *Aegle marmelos*as an alternative or complementary adjunct to conventional anticancer treatments.

## Abbreviation

EAC: Ehrlich’s Ascites Carcinoma
APME: Methanolic Extract
APCE: Chloroform Extract
APHE: n-Hexane Extract
DMSO: Dimethyl Sulfoxide
PBS: Phosphate-Buffered Saline
RH: Relative Humidity
LC50: Median Lethal Concentration
HPLC: High-Performance Liquid Chromatography
i.p.: Intraperitoneal
ICDDR’B: International Centre for Diarrhoeal Disease Research, Bangladesh
SEM: Standard Error of the Mean.

## Acknowledgment

We sincerely express our gratitude to the Department of Pharmacy, Jahangirnagar University, Dhaka, Bangladesh, for providing laboratory facilities and technical support throughout this research. We also extend our appreciation to the Department of Biochemistry and Molecular Biology, University of Rajshahi, for generously providing EAC cells.

